# Monitoring data indicates some annual change in the mammal fauna at Nitmiluk National Park between 2005 and 2018 but a reduction in effort confounds any interpretation

**DOI:** 10.64898/2026.05.21.726992

**Authors:** A.S. Kutt, A. Edwards, H.S. Fraser

## Abstract

**Context:** The decline, extirpation and extinction of Australias mammal fauna is without peer. In recent decades long term monitoring has revealed tragic patterns in iconic locations such as Kakadu National Park.

**Aims:** Past studies have described universal patterns across multiple national parks in the Northern Territory, linking the declines to fire management; however, much of the data in Nitmiluk National Park do not cover the period that encompasses this decline. We examine this data separately to look for patterns with respect to fire history.

**Methods:** We examined standardised mammal survey data collected in Nitmiluk from 2005 to 2018, to look for correlates with year of survey, time since last burnt, and the frequency of early season and late season burns.

**Key results:** Despite subtle difference in methods over the years, we found there were some annual changes in the mammals, but little discernible pattern of change correlated to fire regime (which itself indicated an improvement in seasonality, and the extent of burning overall over this period). Instead, there was a distinct reduction in the survey effort.

**Conclusions:** Our study provides further support for the pitfalls of ecological monitoring losing focus, being decoupled from management actions, and allowing ill-conceived changes in design. We recommend that future monitoring at Nitmiluk should focus on key management questions (i.e., fire), best methods for target taxa (i.e. camera traps), more regular and flexible sampling linked to a conceptual model, and integration and co-design with the local land managers, rangers and Traditional Owners.

**Short Summary:** Re-examination of the mammal data collected in monitoring sites at Nitmiluk National Park between 2005 to 2018 indicated little pattern with fire regime over time, and instead indicated that a reduction in effort, had reduced the ability to find pattern With sparse and declining mammal population in northern Australia, future monitoring needs to be focused on key management questions.

## Introduction

The decline, extirpation and extinction of Australian mammals has been well documented (Woinarski *et al*. 2015). Since the first wave of loss of a suite of unique medium sized mammals, there has been increasing concern that a second wave of disappearance has commenced in the remaining mammals, and particularly in the small dasyurids and rodents (Fisher *et al*. 2014; Lawes *et al*. 2015a). The evidence is stark for locations such as the Top End of the Northern Territory (Woinarski *et al*. 2010), though there is not the equivalent long term data from north-eastern Australia, aside from data indicating the abundance of mammals, collected via similar standardised sampling methods, is equally low (Perry *et al*. 2015). Regardless, there is substantiation that key threats are universal in northern Australia and that changes in vegetation cover and composition, caused by grazing by domestic stock and fire, feral cat predation, and poisoning and predation by the introduced Cane Toad, are all culpable (Kutt 2012; Legge *et al*. 2019; von Takach *et al*. 2022). The relative and interacting effects, and the means to manage these threats in mainland landscapes outside fenced enclosures and islands, is complex and not conceptually straightforward (Doody *et al*. 2024; Radford *et al*. 2020).

In the Northern Territory long term monitoring has allowed the recent changes in mammal populations to be clearly and distressingly reported (Woinarski *et al*. 2001). These data have continued to be collected and published, but most recently as a combined data set across multiple parks (Einoder *et al*. 2023), masking the individual patterns in each separate reserve. Furthermore, the mammal fauna is not the same across these national parks (i.e., more species and endemics at Kakadu compared to sites further south and west, and an unequal time span of sampling across the Parks), which undermines some elements of the data use and interpretation. This, and the lack of location specific analysis of the patterns of change, linked to management, creates some confusion regarding the correlation with key threats such as fire. For example, there is historical evidence of poor fire management at some parks (Edwards *et al*. 2001) and subsequent research regarding vegetation patterns (Russell-Smith *et al*. 2009; Russell-Smith and Edwards 2006), that has resulted in fire and carbon programs to arrest and change inappropriate fire management (Russell-Smith *et al*. 2015). Matching site-specific wildlife data on the benefits or otherwise of these programs, is limited.

In this study we examined mammal data collected via fauna surveys in Nitmiluk National Park, Northern Territory from 2005 to 2018. In this case, we investigated two simple questions, namely, (i) what is the change in species richness and abundance over the years of survey at Nitmiluk National Park? (ii) does fire history or rainfall predict the patterns in the mammal data? Undertaking more nuanced location specific analysis of data, is of value in both unmasking more accurate patterns, or declines, and then focussing and improving management on conservation reserves.

## Methods

### Study Site

The data were collected in Nitmiluk National Park (hereafter NNP): in the Northern Territory. NNP is in the Katherine region, approximately 320km south of Darwin (14.0873° S, 132.4971° E). The park is approximately 290,000 ha in size, includes a portion of the Arnhem Plateau and shares a south-western boundary with Kakadu National Park (Fig. 1). The vegetation is predominantly open woodlands and forests, includes sandstone escarpment woodlands and heath, monsoon vine-thicket and other closed forests (i.e., riparian), grasslands and wetlands.

**Figure 1.**
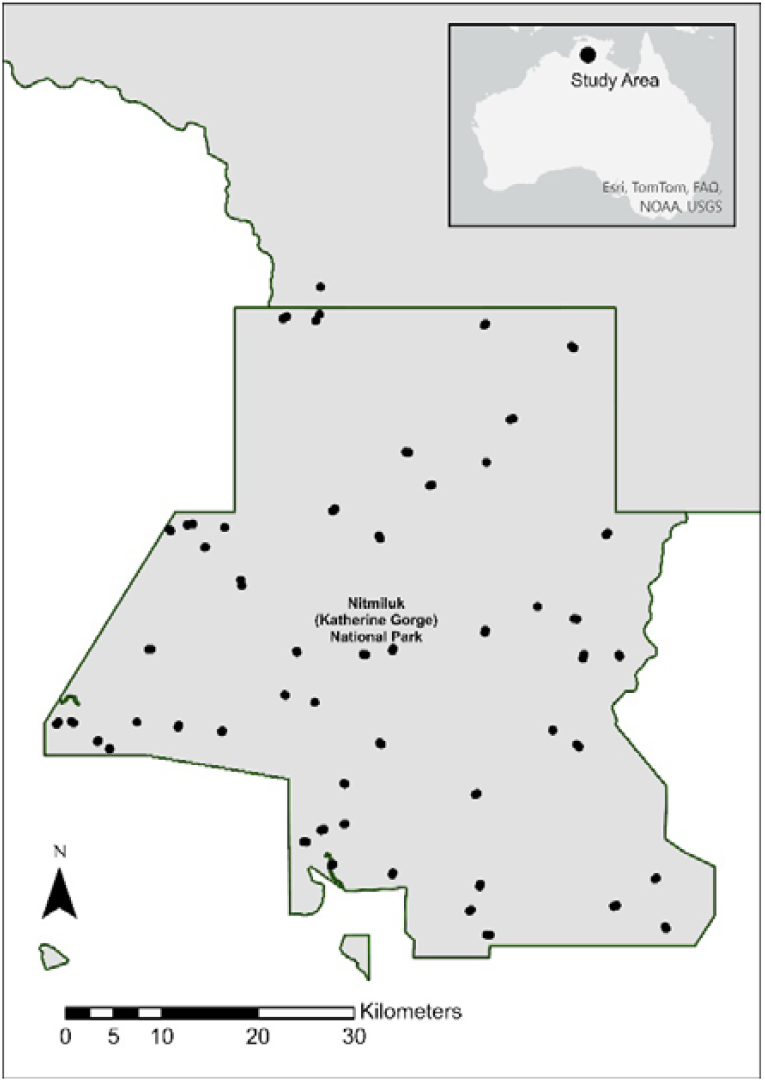
The location of Nitmiluk National Park, Northern Territory, Australia. The black circles are all the monitoring sites sampled over the years of survey.

**Figure 2.**
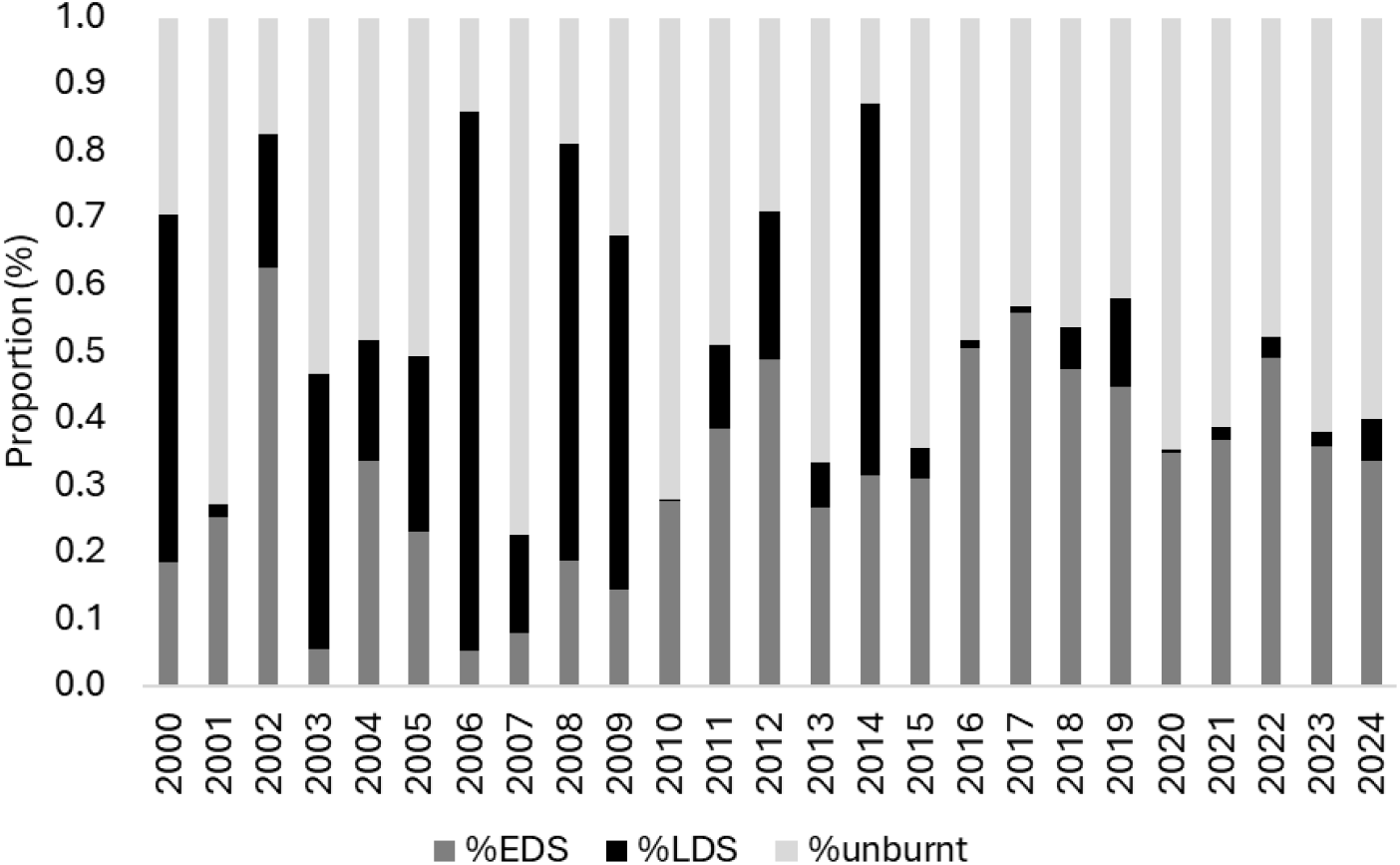
The percentage distribution of early dry season (%EDS - yellow), late dry season (LDS% - red) and total area unburnt (%unburnt - green) at Nitmiluk National Park from 2000-2018.

**Fig. 3.**
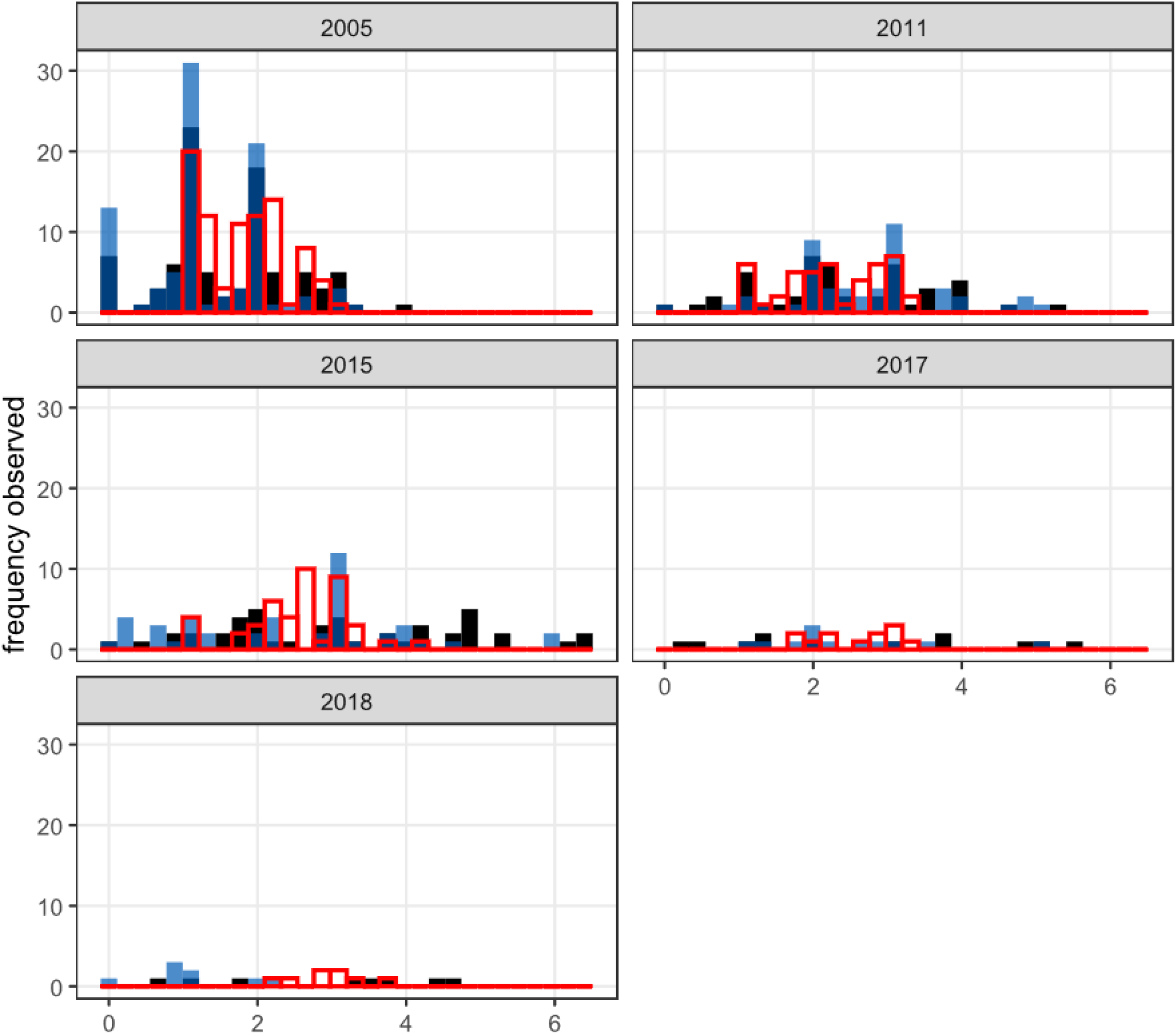
The number of sample sites each year of survey, for time since last burn (empty bars with red outline), the frequency (number of times in previous 10 years) of early dry season burns (solid black bars) and late dry season burns (solid blue bars).

### Surveys

We examined five surveys conducted at NNP between 2005 and 2018. The methods used were a systematic site approach used throughout the Northern Territory (Einoder *et al*. 2020; Woinarski *et al*. 2010). In this study we only investigated the mammal data, though reptile, amphibian and bird data were also collected.

The standard effort for the surveys varied across the sites and years, but consisted of Elliott, cage and pitfall traps, and diurnal and nocturnal active searching. Funnel traps were used in the later year surveys, but this method does not trap mammals. Over the five surveys we examined, the number of survey nights ranged from two to four, the number of Elliott traps ranged from 16-30, the number of cages set ranged from 4-8, and the number of pitfall traps ranged from 0-4. Nocturnal active searching was consistent across all surveys (two 20- minute searches), though in the 2017 and 2018 surveys these were four 10-minute searches (Supplementary Table 1).

We standardised our data to represent species richness or abundance per 100 trap nights. The total trap effort was the sum of the Elliott, cage and pitfall traps set at each site multiplied by the number of nights of survey. Trap nights ranged from 72-112. The site species richness or abundance was divided by this figure and multiplied by 100. For example, a site sampled over 2 nights by 20 Elliotts, 4 cages and 4 pitfall traps, represents 56 trap nights. If a species scored an abundance of 1 for that site, then the relative abundance for 100 trap nights, is 1/(2*28) *100 or 1.78. As the active search effort was consistent across all surveys, these data were included in the abundance for a species at a site before the “per 100 trap night” standardisation; active searching generally recorded arboreal nocturnal mammals, of which there were very few recorded.

We only considered small native mammal species (<5.5 kg) of which there were 14 recorded, representing one monotreme, nine marsupials and four rodents (Table 2). From the survey data we derived the total records per species per site (i.e., relative abundance) and abundance and species richness per site for all mammals, and then for marsupials (excluding monotremes) and rodents. There is a suite of mammals, recorded from Kakadu National Park, and included in cross-park analysis of temporal changes and management (Einoder *et al*. 2023) that have not been recorded from the monitoring sites at Nitmiluk, and these are; Rakali *Hydromys chrysogaster*, Rock ringtail possum *Petropseudes dahli*, Northern Brush-tailed Phascogale *Phascogale pirata*, Northern Quoll *Dasyurus hallucatus*, Northern Short-tailed Mouse *Leggadina forresti*, Pale Field-rat *Rattus tunneyi*, Kakadu Pebble-mouse *Pseudomys calabyi*, Fawn Antechinus *Antechinus bellus*, Dusky Rat *Rattus colleti*, Brush-tailed Rabbit-rat *Conilurus penicillatus* and Arnhem Rock Rat *Zyzomys maini*.

### Environmental data

The fire data were produced by North Australia and Rangelands Fire Information (NAFI) at Charles Darwin University, derived from satellite imagery sourced from the Moderate Resolution Imaging Spectroradiometer (MODIS) using the Red and Near Infrared bands with a 250 x 250 m pixel resolution. The fire metrics, time since last burnt (TSLB), frequency of early season burns (FRQE) and the frequency of late season burns (FRQL) were calculated using standard GIS techniques covering the 10 years prior to each of the survey years (ESRI Inc. 2024). Early dry season commences usually in May, at the end of the annual wet season (November to April), through to July, and the late dry season occurs from August to October, or later, depending on the commencement of the following wet season. The fire metrics were attributed to each site using the mean value of a 3 x 3-pixel window centred over the survey location.

Monthly (and therefore annual) rainfall data were extracted from the Bureau of Meteorology monthly gridded rainfall dataset (http://www.bom.gov.au/). The grids are generated using statistical or optimal interpolation which is basically a form of least squares regression applied in two dimensions (Daley 1993). See Evans *et al*. (2020) for full details of the analysis approach. Rainfall is the total for the 12-months prior to the surveys.

### Analysis

We began by plotting the mean and 95% confidence intervals of each of the environmental variables (FRQE, FRQL, TSLB, and rainfall, Fig. 5), and each of the mammal variables (total, rodent, and marsupial richness and abundance, Fig. 4) by year to examine general trends in the data.

**Fig. 4.**
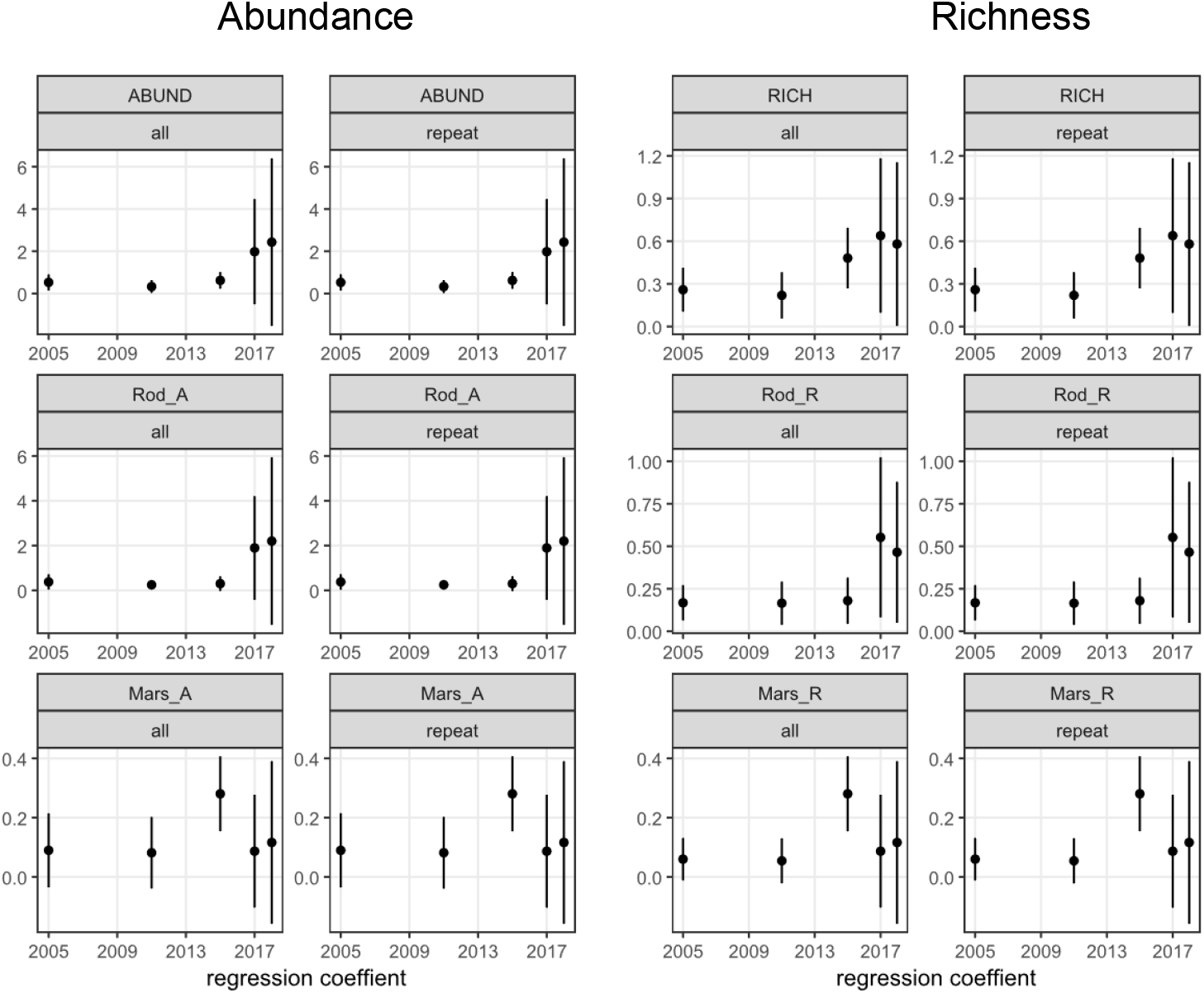
The mean and 95% confidence intervals for all native mammal, rodent and marsupial abundance and species richness for the full data set across the years (2005 to 2017) and then the sites that were repeat sampled three times. **all** = full data set and **repeat** = repeat sampled three times.

**Fig. 5.**
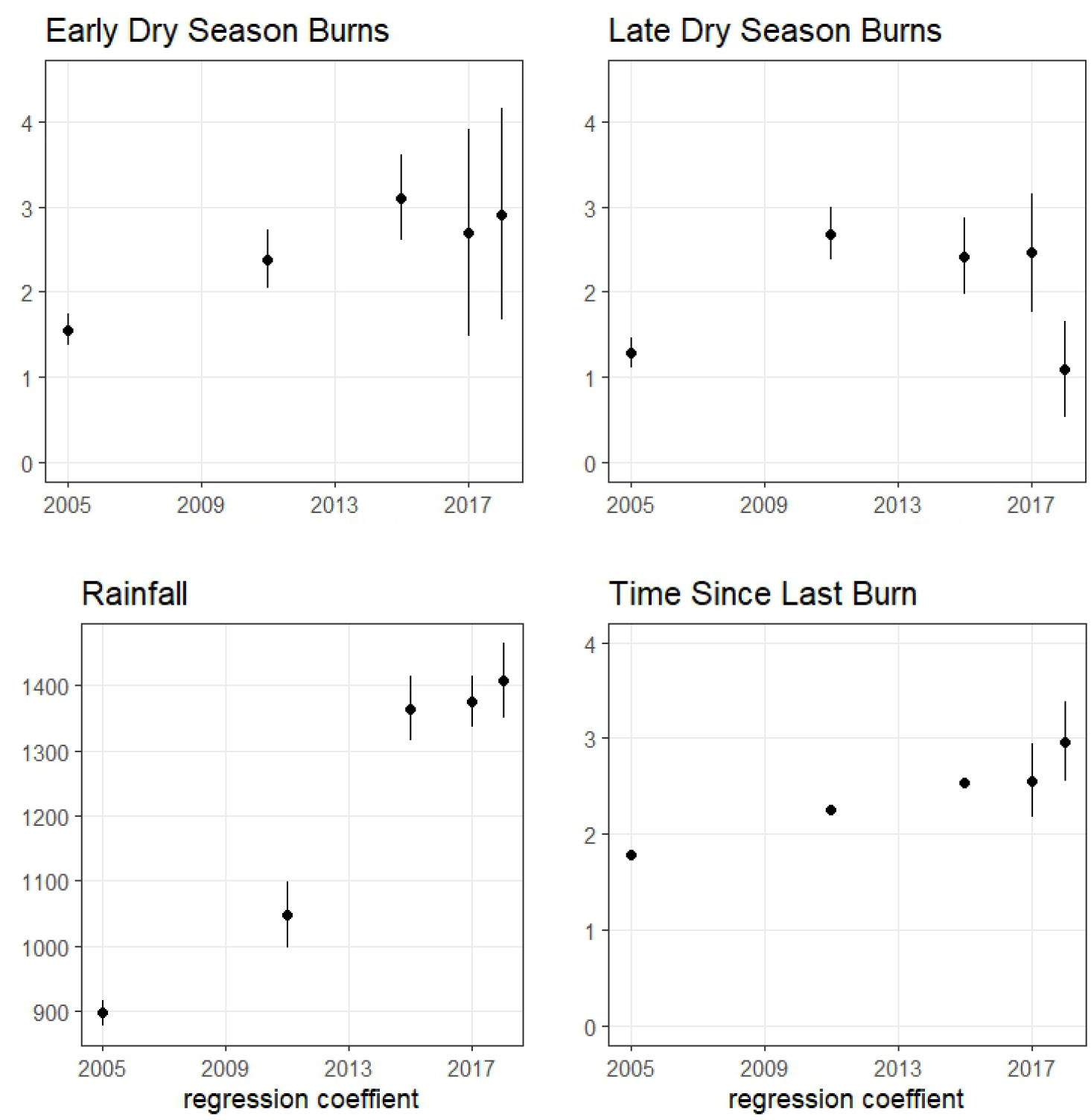
The site means and upper and lower 95% confidence intervals for the four environmental variables for each year of survey.

We used simple generalised linear models in R version 4.4.3 (R Core Team 2024) to describe the relationship between year and richness and abundance of all mammals (Table 1), marsupials and rodents (Supplementary Material Table 2) and species if recorded in 10 or more sites (Supplementary Material Table 3). We did this for the whole dataset. As the number of sites sampled reduced drastically over time, we repeated this analysis, only including sites that had been surveyed in more than 3 years to reduce bias in the estimated trend through time.

**Table 1.** The generalised linear model results for the relationship between abundance and richness with year for full dataset and for dataset subset to sites which were surveyed in at least 3 years. Then excluding 2005 as there’s no change in richness or abundance until 2011.

| Sites | Years | Variable | Term | Estimate | S.E. | Statistic | P value | 95% CI low | 95% CI high |
| --- | --- | --- | --- | --- | --- | --- | --- | --- | --- |
| All | All | Richness | (Intercept) | -137.55 | 52.34 | -2.63 | 0.01 | -241.79 | -35.84 |
| All | All | Richness | YEAR | 0.07 | 0.03 | 2.61 | 0.01 | 0.02 | 0.12 |
| All | All | Abundance | (Intercept) | -175.00 | 37.76 | -4.63 | 0.00 | -250.08 | -101.79 |
| All | All | Abundance | YEAR | 0.09 | 0.02 | 4.63 | 0.00 | 0.05 | 0.12 |
| All | excluding 2005 | Richness | (Intercept) | -324.04 | 126.98 | -2.55 | 0.01 | -579.93 | -79.81 |
| All | excluding 2005 | Richness | YEAR | 0.16 | 0.06 | 2.55 | 0.01 | 0.04 | 0.29 |
| All | excluding 2005 | Abundance | (Intercept) | -629.20 | 102.10 | -6.16 | 0.00 | -835.93 | -434.83 |
| All | excluding 2005 | Abundance | YEAR | 0.31 | 0.05 | 6.16 | 0.00 | 0.22 | 0.41 |
| 3+ repeat | All | Richness | (Intercept) | -104.95 | 65.95 | -1.59 | 0.11 | -238.02 | 21.67 |
| 3+ repeat | All | Richness | YEAR | 0.05 | 0.03 | 1.58 | 0.11 | -0.01 | 0.12 |
| 3+ repeat | All | Abundance | (Intercept) | -173.67 | 49.04 | -3.54 | 0.00 | -272.34 | -79.67 |
| 3+ repeat | All | Abundance | YEAR | 0.09 | 0.02 | 3.54 | 0.00 | 0.04 | 0.14 |
| 3+ repeat | excluding 2005 | Richness | (Intercept) | -382.01 | 146.88 | -2.60 | 0.01 | -678.59 | -99.96 |
| 3+ repeat | excluding 2005 | Richness | YEAR | 0.19 | 0.07 | 2.60 | 0.01 | 0.05 | 0.34 |
| 3+ repeat | excluding 2005 | Abundance | (Intercept) | -747.83 | 118.43 | -6.31 | 0.00 | -987.50 | -522.53 |
| 3+ repeat | excluding 2005 | Abundance | YEAR | 0.37 | 0.06 | 6.32 | 0.00 | 0.26 | 0.49 |

**Table 2.** The mean (and standard error) for the environmental variables and the mammal groups and species for each year.

| Variable / group | Total sites | 2005 | 2011 | 2015 | 2017 | 2018 |
| --- | --- | --- | --- | --- | --- | --- |
| Number of sites | 95 | 88 | 46 | 45 | 12 | 8 |
| Time since last burnt |  | 1.78 (0.06) | 2.26 (0.10) | 2.53 (0.10) | 2.56 (0.18) | 2.96 (0.18) |
| Frequency early dry season |  | 1.55 (0.09) | 2.38 (0.17) | 3.11 (0.25) | 2.69 (0.55) | 2.91 (0.52) |
| Frequency late dry season |  | 1.28 (0.09) | 2.68 (0.16) | 2.42 (0.22) | 2.46 (0.31) | 1.09 (0.24) |
| Rainfall preceding 12 months |  | 897.90 (9.89) | 1048.30 (25.19) | 1364.20 (24.29) | 1375.33 (17.93) | 1408.12 (24.45) |
| Mammal species richness |  | 0.26 (0.08) | 0.22 (0.08) | 0.48 (0.11) | 0.64 (0.25) | 0.58 (0.24) |
| Mammal abundance |  | 0.53 (0.19) | 0.33 (0.15) | 0.62 (0.20) | 1.98 (1.13) | 2.43 (1.68) |
| Total species recorded |  | 9 | 8 | 12 | 8 | 8 |
| Rodent abundance |  | 0.38 (0.18) | 0.25 (0.11) | 0.30 (0.17) | 1.89 (1.05) | 2.20 (1.58) |
| Rodent species richness |  | 0.17 (0.05) | 0.17 (0.06) | 0.18 (0.07) | 0.55 (0.21) | 0.46 (0.18) |
| Marsupial abundance |  | 0.09 (0.06) | 0.08 (0.06) | 0.28 (0.06) | 0.09 (0.09) | 0.12 (0.12) |
| Marsupial species richness |  | 0.06 (0.04) | 0.05 (0.04) | 0.28 (0.06) | 0.09 (0.09) | 0.12 (0.12) |
| **Monotreme** |  |  |  |  |  |  |
| *Tachyglossus aculeatus* | 4 | 0.03 (0.02) |  | 0.02 (0.02) |  | 0.12 (0.12) |
| **Marsupials** |  |  |  |  |  |  |
| *Isodon macrourus* | 2 |  |  | 0.02 (0.02) |  | 0.12 (0.12) |
| *Petaurus ariel* | 4 | 0.03 (0.03) | 0.05 (0.05) | 0.04 (0.03) |  |  |
| *Planigale maculata* | 7 | 0.02 (0.01) |  | 0.12 (0.05) |  |  |
| *Pseudantechinus bilarni* | 1 | 0.03 (0.03) |  |  |  |  |
| *Pseudomys nanus* | 1 |  | 0.03 (0.03) |  |  |  |
| *Sminthopsis bulteri* | 1 |  |  | 0.02 (0.02) |  |  |
| *Sminthopsis nitela* | 4 | 0.02 (0.01) |  | 0.04 (0.03) | 0.09 (0.09) |  |
| *Trichosurus vulpecula* | 2 |  |  | 0.04 (0.03) |  |  |
| *Petrogale wilkinsi* | 3 | 0.02 (0.02) |  | 0.02 (0.02) |  |  |
| **Rodents** |  |  |  |  |  |  |
| *Melomys burtoni* | 15 | 0.30 (0.17) | 0.16 (0.10) | 0.12 (0.10) | 1.66 (1.05) | 1.97 (1.61) |
| *Mesembriomys gouldii* | 2 |  |  | 0.04 (0.03) |  |  |
| *Pseudomys delicatulus* | 6 | 0.03 (0.02) |  | 0.04 (0.03) | 0.15 (0.10) |  |
| *Zyzomys argurus* | 10 | 0.05 (0.03) | 0.08 (0.05) | 0.10 (0.07) | 0.08 (0.08) | 0.23 (0.23) |

We used hierarchical generalized linear mixed models to investigate the influence of FRQE, FRQL, TSLB, and rainfall, with a random effect of year, for mammal groups and species as above. We did this for both the full dataset, and the dataset which only includes repeat-sampled sites. All these models encountered failure to converge errors. Examination of the data (Fig. 3) reveals that there is an insufficient number of sites in later years, and therefore there is insufficient variation in the fire variables to support these analyses.

We tabulated the mean (and S.E.) of abundance and species richness for the three mammal groups, and the mean (and S.E.) of abundance of all mammal species recorded, as well as the fire variables and rainfall (Table 2).

## Results

We found an increase in mammal richness and abundance with time (Table 2), which was robustly present regardless of which subset of the data we took. There appears to be an increase in rainfall, time since last burn, and early dry season fires with time, though the relationship between late dry season fires and time is less clear (Fig. 5). Any or all these factors may have contributed to the observed increase in mammal richness and abundance, but the lack of data did not support analysis of the relationship between mammals and these environmental variables.

There were 95 unique sites sampled over the survey years. The number of sites sampled each year reduced over time (86 in 2005, 46 in 2011, 45 in 2015, 12 in 2017 and 8 in 2018). The number of repeat samples were also low, with 41 sampled once, 12 sampled twice, 34 three times, 8 four times, and 0 five times.

The early and late dry season burn pattern at the mammal monitoring sites (Fig. 5) do not match the broader trends in burning across the park (Fig. 2). At the monitoring sites, the frequency of late and early dry season fires is reasonably similar (Fig. 5), while at a park scale, early dry season fires are markedly more common (Fig. 2). The area of unburnt, early dry season (EDS) and late dry season (LDS) fires for the entire national park varied quite considerably from 2000 to 2024 (Fig. 2). Considering the years of survey and the preceding extent of burning, from 2000-2005, the areas were 45% unburnt, 28% early dry season burnt, and 26% late dry season burnt. From 2006-2011, the areas were 44% unburnt, early dry season burnt 18% and 37% late dry season burnt. From 2012-2018, these totals were 44%, 31% and 13% and 2019-2024, 56%, 39% and 5% respectively.

## Discussion

In this study we examined the changes in mammals sampled systematically at Nitmiliuk National Park between 2005 and 2018 and investigated if any changes in abundance and species richness could be correlated to changing fire management patterns over the years of survey. Though there was a hint of increases in some species and the total mammal pool, we could not attribute any of the changes in the data to fire history, even though there seemed to be an improvement in the overall pattern and an increase in the extent of early dry season burns. Though we recognise that there were subtle changes in the survey methods over the years, we believe the differences were not sufficient to prevent comparison of the data sets. However, what was apparent, was a marked reduction in survey effort, as measured by the number of sites sampled year on year. There was an equally marked reduction in the range and variation of fire.

The original intent of the Nitmiluk standardised surveys, commenced in 2005, were to provide baseline and subsequent fauna data for management, with sites linked to long term fire and vegetation dynamic research (Edwards *et al*. 2001) as well as to describe the biogeographic variation and conservation benefits of the large national parks in the region (Woinarski *et al*. 2013). As some troubling changes were beginning to manifest in mammal populations (Woinarski *et al*. 2001), the focus of the data analysis shifted to correlating declines to threats such as unmanaged fire (Lawes *et al*. 2015b). Subsequent monitoring changed the sampling approach to increase the site-based effort (to increase the power of the sampling) which also then decreased the total number of sites surveyed across each of the parks (Einoder *et al*. 2020) and grouped sites across different geographic locations and different time frames (i.e., Three Parks Fire Plot Network, Einoder *et al*. 2023). However, the total mammal pool in Kakadu National Park represents 25 mammal species (<5 kg) from surveys between 1996-2019, Litchfield National Park, 19 between1995-2016 and Nitmiluk National Park only 14 over 2005-2018. This inequality in the base mammal fauna (i.e., some species are absent from each park) undermines the confidence in recent analysis regarding changes in the mammal populations and the correlative explanations (i.e., fire pattern). Our analysis of the Nitmiluk data indicates both the site number and species composition are too few to derive any meaningful pattern; as such, there is no evidence of mammal decline from the data collected between 2005 and 2018 and we cannot conclude if mammals declined at Nitmiluk prior to 2005, though it is very likely they have.

Our examination of the Nitmiluk data (in which the initial sampling coincided with, or post-dated, the Kakadu declines), hints slightly of some annual change in mammals, driven by two species (*Melomys burtoni* and *Zyzomys argurus*) and maybe by fire pattern. Other more focussed analysis of mammal changes in northern Australia, provide data that indicates there are complex interacting reasons for shifts in mammal population, such as the nexus between increased fire frequency, increased feral predation and decline in habitat quality (i.e., shelter and food resources) for species (Stobo-Wilson *et al*. 2025). Some species, such as *Melomys burtoni* are disturbance tolerant, and can thrive in agricultural and partially cleared environments (Dyer *et al*. 2011). In this study we have only examined a very limited set of predictive variables, focussed on fire as this variable has consistently been correlated with mammal change (Lawes *et al*. 2015b). However more sophisticated experimental research that attempts to unpick reasons for mammal changes due to fire management, also point to survival, recruitment and diet (Griffiths and Brook 2015) and idiosyncratic responses to differences in vegetation structure and composition (Firth *et al*. 2006).

In this study, we originally sought to examine if changes in landscape scale fire management (i.e., across Nitmiluk) could be correlated to changes in mammal pattern, given there are a number of existing analyses that suggest that fire pattern has a relationship with mammal abundance and decline (Einoder *et al*. 2023). Instead, our results indicated that changes in monitoring, to strengthen aspects such as power, and articulate more universal (i.e., multi-park) inferences for management, in fact reduced the ability to provide any meaningful insight to a very large conservation area (290,000 ha) that is managed separately by Traditional Owners and Indigenous Ranger groups (Russell-Smith *et al*. 2015). There is a substantial literature regarding principles and pitfalls of monitoring programs, including for Kakadu National Park (Woinarski 2012), and our examination of the Nitmiluk data, suggests it has fallen foul to a degree of passivity and the absence of key questions or hypothesis, poor experimental design (i.e., progressive reduction in number of sites) and the lack of evolution in the design to shift to, for example, a clear focus on fire management, that integrates with the on-ground fire management programs (Lindenmayer and Likens 2010). Our analysis suggests several recommendations for future monitoring at Nitmiluk, that can incorporate the existing site network (including the sites discontinued from sampling), namely (i) develop a conceptual model to underpin future monitoring, (ii) review the past, current and proposed fire management to carefully stratify sites to examine variation across the landscape and fire strategy, (iii) a focus on methods such as camera trapping, to provide a more cost-effective, data and site rich monitoring array, (iv) more regular sampling and review of the data, and (v) co-design of future monitoring, to integrate with Traditional Owner aspirations and knowledge needs. Consistent long-term monitoring is critical to investigate trends and changes in key taxa over time, though in the data we examined here, we conclude that aspirations for perfection should not be the enemy of good monitoring (Lindenmayer and Likens 2010).

## Acknowledgements

We acknowledge and pay our respects to the Jawoyn people who jointly manage Nitmiluk National Park. The surveys were conducted over the years, by staff and volunteers from the Flora and Fauna Division, Department of Lands, Planning and Environment. All surveys were undertaken under scientific research permits, and animal ethics approvals, as prescribed by the Territory Parks and Wildlife Conservation Act.

