## Supplemental Tables for "Monitoring data indicates some annual change in the mammal fauna at Nitmiluk National Park between 2005 and 2018 but a reduction in effort confounds any interpretation"

**Supplementary Material Table 1.** The survey effort per year including the number of sites sampled. Some effort includes a range of values, and the number in parenthesis is the mean.

| **Year** | **Sites** | **Survey nights** | **Elliot traps** | **Cage traps** | **Pitfall traps** | **Active searches (diurnal)** | **Active searches (nocturnal)** |
| --- | --- | --- | --- | --- | --- | --- | --- |
| 2005 | 88 | 2.0 | 30.0 (29.8) | 5.0-6.0 (5.9) | 0.0-2.0 (1.9) | 3.0 x 20 minutes | 2.0 x 20 minutes |
| 2011 | 46 | 2.0 | 30.0 | 6.0 | 2.0-4.0 (3.7) | 3.0 x 20 minutes | 2.0 x 20 minutes |
| 2015 | 45 | 4.0 | 16.0 | 7.0-8.0 (7.96) | 3.0-4.0 (3.9) | 3.0 x 20 minutes | 2.0 x 20 minutes |
| 2017 | 12 | 3.0-4.0 (3.9) | 16.0 | 8.0 | 0.0-3.0 (2.7) | 4.0-6.0 (5.8) x 10 minutes | 3.0-4.0 (3.8) x 10 minutes |
| 2018 | 8 | 4.0 | 16.0 | 8.0 | 3.0 | 6.0 x 10 minutes | 4.0 x 10 minutes |

**Supplementary Material Table 2.** The hierarchical regression models for mammal groups and Year, indicating the group, predictor, the estimate regression coefficients, standard errors, the test statistic and p value, and the upper and lower 95% confidence intervals for each predictor, and R2 values for the Regression models were undertaken for all samples, and then sites that were resampled at least three times.

| **Groups** | **Term** | **Estimate** | **S.E.** | **Statistic** | **P value** | **95% CI low** | **95% CI high** | **R^2^** |
| --- | --- | --- | --- | --- | --- | --- | --- | --- |
| **All samples** |  |  |  |  |  |  |  |  |
| Mammal species richness | (Intercept) | -132.711 | 62.643 | -2.119 | 0.034 | -255.131 | -13.220 | 0.007 |
|  | YEAR | 0.065 | 0.031 | 2.101 | 0.036 | 0.006 | 0.126 | 0.007 |
| Rodent species richness | (Intercept) | -116.322 | 74.468 | -1.562 | 0.118 | -261.837 | 25.804 | 0.009 |
|  | YEAR | 0.057 | 0.037 | 1.541 | 0.123 | -0.014 | 0.129 | 0.009 |
| Marsupial species richness | (Intercept) | -233.374 | 96.935 | -2.408 | 0.016 | -434.793 | -49.282 | 0.020 |
|  | YEAR | 0.115 | 0.048 | 2.386 | 0.017 | 0.023 | 0.215 | 0.020 |
| Mammal abundance | (Intercept) | -139.550 | 81.813 | -1.706 | 0.088 | -286.939 | 3.146 | 0.005 |
|  | YEAR | 0.069 | 0.041 | 1.700 | 0.089 | -0.002 | 0.143 | 0.005 |
| Rodent abundance | (Intercept) | -154.539 | 109.147 | -1.416 | 0.157 | -349.259 | 30.586 | 0.003 |
|  | YEAR | 0.076 | 0.054 | 1.409 | 0.159 | -0.016 | 0.173 | 0.003 |
| Marsupial abundance | (Intercept) | -156.144 | 100.089 | -1.560 | 0.119 | -360.196 | 37.918 | 0.000 |
|  | YEAR | 0.077 | 0.050 | 1.540 | 0.124 | -0.020 | 0.178 | 0.000 |
| **Resampled sites** |  |  |  |  |  |  |  |  |
| Mammal species richness | (Intercept) | -97.071 | 81.153 | -1.196 | 0.232 | -253.370 | 54.287 | 0.001 |
|  | YEAR | 0.048 | 0.040 | 1.185 | 0.236 | -0.027 | 0.125 | 0.001 |
| Rodent species richness | (Intercept) | -87.878 | 100.192 | -0.877 | 0.380 | -284.701 | 99.929 | 0.002 |
|  | YEAR | 0.043 | 0.050 | 0.862 | 0.389 | -0.050 | 0.141 | 0.002 |
| Marsupial species richness | (Intercept) | -219.920 | 115.861 | -1.898 | 0.058 | -464.395 | -4.651 | 0.027 |
|  | YEAR | 0.108 | 0.058 | 1.883 | 0.060 | 0.001 | 0.230 | 0.027 |
| Mammal abundance | (Intercept) | -121.916 | 104.374 | -1.168 | 0.243 | -298.385 | 53.204 | 0.004 |
|  | YEAR | 0.060 | 0.052 | 1.165 | 0.244 | -0.027 | 0.148 | 0.004 |
| Rodent abundance | (Intercept) | -165.265 | 148.792 | -1.111 | 0.267 | -413.957 | 79.193 | 0.003 |
|  | YEAR | 0.082 | 0.074 | 1.106 | 0.269 | -0.040 | 0.206 | 0.003 |
| Marsupial abundance | (Intercept) | -101.346 | 115.522 | -0.877 | 0.380 | -332.107 | 116.952 | 0.000 |
|  | YEAR | 0.050 | 0.057 | 0.862 | 0.388 | -0.059 | 0.164 | 0.000 |

**Supplementary Material Table 3.** The hierarchical regression models for mammal species and Year, indicating the group, predictor, the estimate regression coefficients, standard errors, the test statistic and p value, and the upper and lower 95% confidence intervals for each predictor, and R^2^ value. Regression models were undertaken for all samples, and then sites that were resampled at least three times.

| **Species** | **Term** | **Estimate** | **S.E.** | **Statistic** | **P value** | **95% CI low** | **95% CI high** | **R^2^** |
| --- | --- | --- | --- | --- | --- | --- | --- | --- |
| **All samples** |  |  |  |  |  |  |  |  |
| *Tachyglossus aculeatus* | (Intercept) | 13.751 | 223.905 | 0.061 | 0.951 | -409.652 | 468.665 | 0.000 |
|  | YEAR | -0.009 | 0.111 | -0.078 | 0.938 | -0.235 | 0.202 | 0.000 |
| *Zyzomys argurus* | (Intercept) | -173.970 | 154.466 | -1.126 | 0.260 | -491.207 | 121.176 | 0.003 |
|  | YEAR | 0.085 | 0.077 | 1.110 | 0.267 | -0.062 | 0.243 | 0.003 |
| *Pseudomys delicatulus* | (Intercept) | -130.934 | 175.281 | -0.747 | 0.455 | -496.773 | 209.256 | 0.001 |
|  | YEAR | 0.063 | 0.087 | 0.728 | 0.467 | -0.106 | 0.245 | 0.001 |
| *Melomys burtoni* | (Intercept) | -147.537 | 175.711 | -0.840 | 0.401 | -466.779 | 142.070 | 0.002 |
|  | YEAR | 0.073 | 0.087 | 0.834 | 0.404 | -0.071 | 0.232 | 0.002 |
| *Planigale maculata* | (Intercept) | -301.644 | 182.751 | -1.651 | 0.099 | -713.262 | 31.395 | 0.013 |
|  | YEAR | 0.148 | 0.091 | 1.633 | 0.102 | -0.017 | 0.353 | 0.013 |
| *Sminthopsis bulteri* | (Intercept) | -224.772 | 223.982 | -1.004 | 0.316 | -733.962 | 195.170 | 0.003 |
|  | YEAR | 0.110 | 0.111 | 0.987 | 0.324 | -0.099 | 0.363 | 0.003 |
| *Petaurus ariel* | (Intercept) | 12.629 | 265.076 | 0.048 | 0.962 | -584.642 | 613.558 | 0.000 |
|  | YEAR | -0.008 | 0.132 | -0.060 | 0.952 | -0.307 | 0.289 | 0.000 |
| **Resamples** | **Term** | **Estimate** | **Standard error** | **Statistic** | **P value** | **95% CI low** | **95% CI high** | **R^2^** |
| *Tachyglossus aculeatus* | (Intercept) | 104.781 | 235.514 | 0.445 | 0.656 | -336.961 | 579.635 | 0.001 |
|  | YEAR | -0.054 | 0.117 | -0.459 | 0.646 | -0.290 | 0.166 | 0.001 |
| *Zyzomys argurus* | (Intercept) | -86.278 | 187.499 | -0.460 | 0.645 | -479.404 | 289.792 | 0.001 |
|  | YEAR | 0.042 | 0.093 | 0.447 | 0.655 | -0.145 | 0.237 | 0.001 |
| *Pseudomys delicatulus* | (Intercept) | -946.514 | 621.641 | -1.523 | 0.128 | -2467.451 | -52.042 | 0.021 |
|  | YEAR | 0.468 | 0.308 | 1.518 | 0.129 | 0.024 | 1.222 | 0.021 |
| *Melomys burtoni* | (Intercept) | -153.077 | 255.410 | -0.599 | 0.549 | -581.941 | 261.372 | 0.002 |
|  | YEAR | 0.076 | 0.127 | 0.595 | 0.552 | -0.130 | 0.289 | 0.002 |
| *Planigale maculata* | (Intercept) | -258.503 | 204.938 | -1.261 | 0.207 | -722.855 | 108.549 | 0.014 |
|  | YEAR | 0.127 | 0.102 | 1.247 | 0.212 | -0.055 | 0.358 | 0.014 |
| *Sminthopsis bulteri* | (Intercept) | -1181.289 | 845.907 | -1.396 | 0.163 | -3292.985 | -17.010 | 0.022 |
|  | YEAR | 0.584 | 0.420 | 1.393 | 0.164 | 0.006 | 1.631 | 0.022 |
| *Petaurus ariel* | (Intercept) | 124.990 | 280.444 | 0.446 | 0.656 | -458.472 | 770.870 | 0.000 |
|  | YEAR | -0.064 | 0.139 | -0.456 | 0.648 | -0.385 | 0.227 | 0.000 |
